# Analytical meditation improves physiological well-being in expert practitioners: a study on central and peripheral neurophysiological correlates

**DOI:** 10.64898/2025.12.08.692357

**Authors:** Alejandro Luis Callara, Mohammad Hadi Azarabad, Laura Sebastiani, Ngawang Sherab, Jampa Khechok, Jampa Tsering, Jampa Thakchoe, Jampa Soepa, Nicola Vanello, Enzo Pasquale Scilingo, Alberto Greco, Bruno Neri

## Abstract

Meditation has been long associated with improvements in mental well-being, emotional regulation, and attentional control. Yet, the diversity of meditative techniques and participant expertise has hindered the systematic identification of their neurophysiological correlates supporting these benefits. To address this challenge, we investigated the neurophysiological signatures of concentrative and analytical meditation in 35 experienced Tibetan monk practitioners. EEG, ECG, EDA, and respiration were simultaneously recorded during *Concentrative, Loving-Kindness* and *Emptiness* meditations. Linear-mixed-models revealed significant modulations in autonomic and cortical activity across meditations. Peripheral indices indicated enhanced parasympathetic tone, decreases respiratory rate, and gradual increase in EDA – reflecting a state of relaxed-alertness with concurrent vagal engagement and sustained sympathetic arousal. EEG analyses supported this state showing elevated gamma-band power during analytical meditations. These findings suggest that advanced meditative states foster an adaptive integration of autonomic and cortical responses, supporting the emergence of relaxed-vigilance – a psychophysiological condition associated with well-being and cognitive flexibility.

## Introduction

Meditation has emerged as a deeply beneficial approach to enhancing well-being, moving beyond its traditional contemplative roots to become a widely studied intervention^1,2^. From a psychological and cognitive perspective, regular meditation practice is linked to tangible improvements in managing stress^3^ and anxiety^4^, fostering emotional regulation^5^, and sharpening cognitive capacities such as attention^6^ and self-awareness^7^, although adverse effects are also reported^8^. This is typically achieved through practices that facilitate a transition from reactive, often self-critical, mental states to others characterized by enhanced clarity, calm, and sustained attention. This transition provides a powerful tool to deepen one’s inner exploration and to improve one’s overall well-being^3^.

Despite the documented benefits of meditation, studies seeking to quantify its overall effects face significant challenges due to the great heterogeneity in contemplative practices^9^. In many contemporary contexts, particularly in the West, meditation is taught and practiced in diverse and sometimes loosely structured ways, leading to substantial variability in technique, adherence, and practitioner experience. ^10^. This variability can make it difficult to determine whether observed effects reflect stable features of meditation or simply differences in how individuals engage with these practices. Accordingly, it remains uncertain to what extent a common physiological or psychological signature of meditation exists across such heterogeneous approaches. In this context, studying more homogeneous and traditionally trained cohorts can provide complementary insights. While such samples cannot represent the full diversity of modern meditation practices, they offer a clearer window onto the underlying mechanisms that may support meditation’s effects. By examining practices that are taught consistently and performed by experienced meditators, it becomes possible to identify more stable neurophysiological patterns, which in turn may help clarify the processes that could contribute to well-being across different forms of meditation.

This is the reason why a recent trend has emerged, promoting the collaboration between Western universities and Eastern monastic institutions^11–17^. These latter institutions provide access to uniquely homogeneous communities of practitioners, offering a research setting where scientists can work with meditators sharing practices and related taxonomy established over centuries. Crucially, even the least experienced monks in these monastic settings may possess years of structured, institutionally guided teachings, which far surpasses the training level of the average Western practitioner cohort. This collaboration thus represents a powerful methodological solution, enabling the study of stable, advanced meditative effects, and paving the way for a more reliable characterization of the effects of meditation.

Two major avenues have been pursued in the characterization of the effects of meditation on the mental state both during and after the meditation session: one based on the use of structured questionnaires to assess behavioral and psychological measures, and the other focused on the neurophysiological processes in central and autonomous nervous system whose effects on mental state and, in particular, on well-being, are well established^18^. The former approach, however, may suffer from significant cultural and conceptual differences that can invalidate some of the results. For example, the application of standardized trait mindfulness scales (developed primarily in Western societies) to Eastern populations risks introducing unwanted cultural bias. This bias can stem from cultural and conceptual divergence^19^, as these scales may fail to accurately assess constructs within other traditional contexts different from the Western.

In this light, the use of neurophysiological correlates provides a necessary alternative objective measure^20^. These markers are less prone to subjective and cultural bias, allowing for a more proper characterization of meditation effects. To this end, two main physiological systems are typically studied: the autonomic nervous system (ANS) using measures such as heart-rate-variability (HRV^21^) and electrodermal activity (EDA^22^), and central nervous system (CNS) using measures such as (magneto)electroencephalography (M/EEG), functional near infrared spectroscopy (fNIRS) and functional magnetic resonance imaging (fMRI). These two families of measures provide both an overview of sympathetic (SNS) and parasympathetic (PNS) nervous system activity, such as arousal or vagal tone, as well as information about brain activity and connectivity. By combining these central and peripheral correlates, it is possible to develop a comprehensive, unbiased view of the physiological changes that accompany advanced meditative states, together with their role in health and well-being. Moreover, the availability of modern, non-invasive sensors now makes it possible to obtain a multivariate window on the subject’s neurophysiology during practice, with minimal interference, as well as to carry out measurements in more ecological settings, altering as little as possible the conditions under which the subject normally performs their practice.

Here, we present a study investigating the central and peripheral neural correlates of concentrative and analytical meditation in a unique and invaluable community: the Monks and Geshes of Sera Jey, one of the oldest Tibetan monastic universities. This study stems from a collaboration between the University of Pisa (Italy) and the Monastic University of Sera Jey (Karnataka, India) and aims to provide a physiological perspective to the characterization of different types of meditation. We recorded central (EEG) and peripheral physiological activity (HRV, EDA, respiration) from 35 experienced meditators performing a three-phase meditation protocol. This protocol included *Concentrative* as well as two distinct forms of *Analytical* meditation: *Loving-Kindness* and *Emptiness* meditation. By analyzing the differential changes in central and peripheral neural correlates across these states, our primary goal is to precisely delineate the neurophysiological signatures of these types of meditation and their role in well-being.

## Methods

### Participants

A total of 35 healthy male participants were recruited (age: 46.1 ± 4.67 years) for this study. All subjects were Geshes and monks of the Sera Jey Monastery. The study was approved by the Committee on Bioethics of the University of Pisa (Review No. 26/2023). The study procedures were conducted in accordance with local legislation and institutional requirements. Written informed consent was obtained from all participants prior to their involvement in the study. All experimental procedures adhered to the principles of Declaration of Helsinki.

### Experimental Design and Protocol

In Figure 1, a summary of the experimental protocol and setup is reported. Subjects were welcomed at the Science Centre of the Sera Jey Monastic University. Prior to the experiment, subjects filled in a demographic and experiential questionnaire, reporting their level of experience, type of meditation usually practiced and daily practice time, as well as some demographics^11^. Then, the experiment began and subjects were asked to perform the experimental protocol reported in Figure 1 (bottom panel). Briefly, after initial 2-minutes eyes-open (EO) and 5-minutes eyes-closed (EC) resting conditions, subjects performed a 3-phase meditation session, comprising concentrative (C, 15 minutes), loving-kindness (LK, 15 minutes) and emptiness (E, 15 minutes) meditation, the latter two being types of analytical meditation. The order of phases was fixed for all participants in accordance with the procedure commonly adopted in Tibetan Buddhist meditation practice, which provides for an initial part of the session devoted to concentrative meditation, aimed at relaxing the body and focusing attention on a neutral object that generates neither attachment nor aversion, until conceptual thought subsides, before performing analytical meditation within the same session. Finally, subjects performed a second EC and EO sessions of two minutes each. The beginning of each session was communicated verbally by the experimenter, and an event marker was placed in the EEG acquisition to identify the beginning of each session.

**Figure 1.**
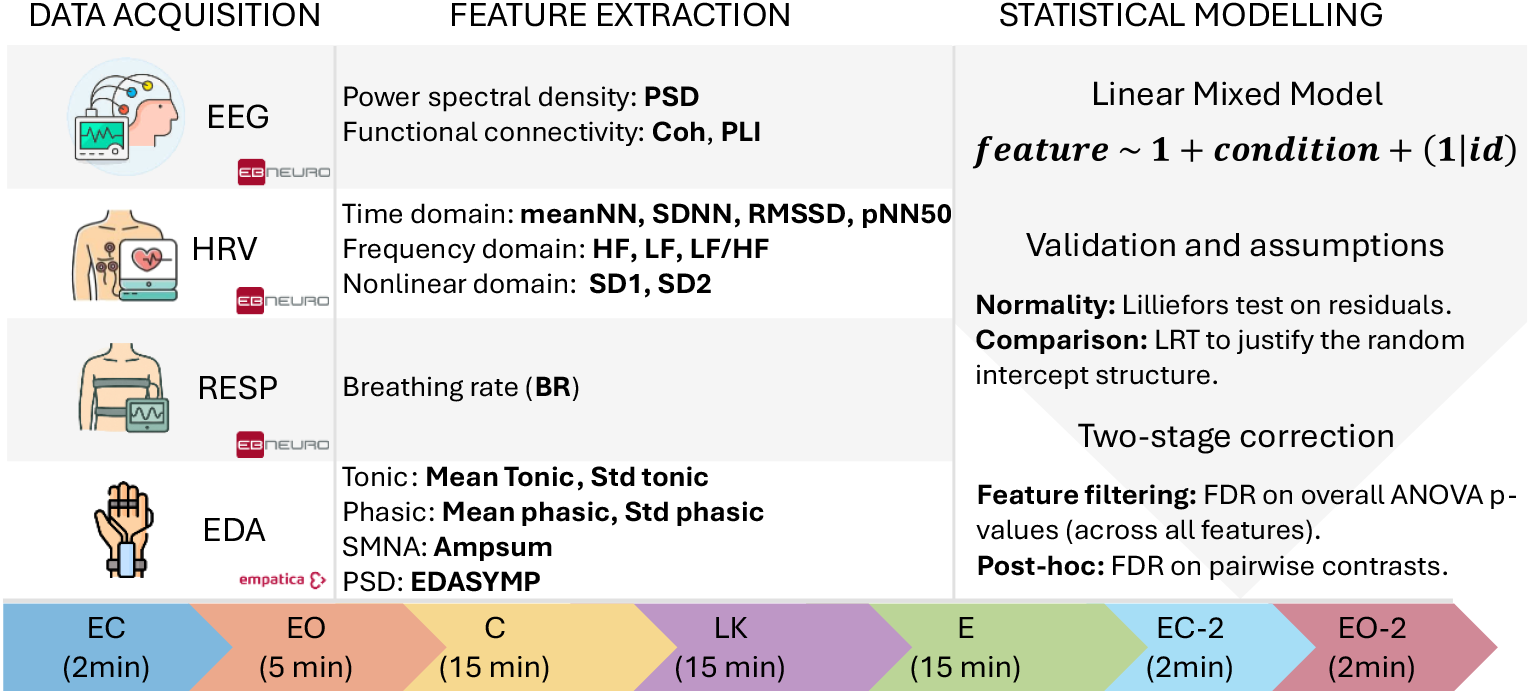
Top (from left to right): Data acquisition, feature extraction, statistical modelling. Bottom: experimental timeline.

**Figure 2.**
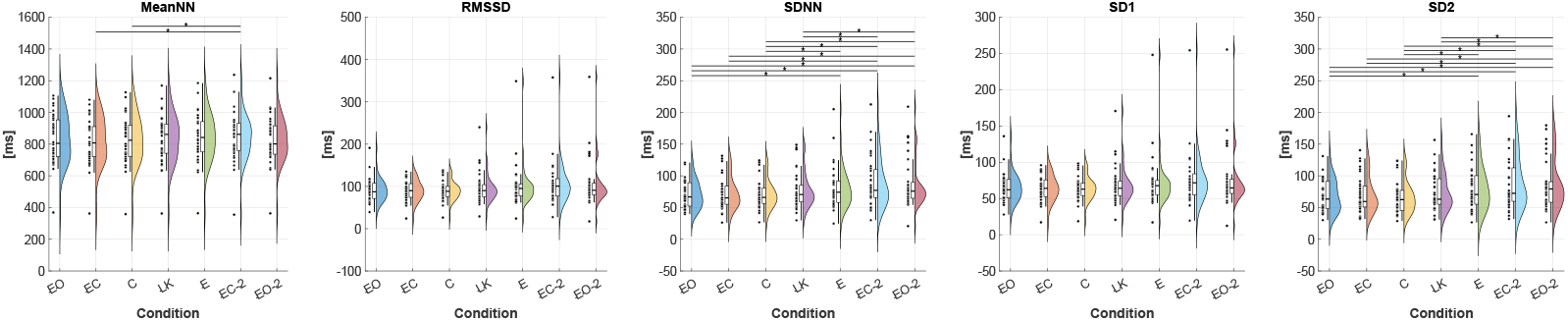
HRV changes. Post-hoc comparisons that were significant (p<0.05) are marked with an asterisk *.

### Description of meditative practices

#### Concentrative meditation^11^

In this practice, the meditator focuses the mind undistractedly on a single point, such as a visualized image (e.g., the Buddha), the natural flow of the breath, or the recitation of a mantra, without analytical elaboration. This meditation develops the ability to focus and serves as the basis for other types of practice. The intention is to cultivate a state of wakefulness that is neither fluctuating nor dull, working to pacify the two main obstacles to concentration: laxity and excitement, to achieve the “factor of clarity” and “factor of stability.”

#### Loving kindness^23^

This practice involves two main approaches. The first is to recollect the kindness of a specific person (often the mother) and focus on the feeling of affection and concern for that person’s wellbeing. The meditator then gradually extends this same sense of loving-kindness to other beings – first to friends and family, then to strangers, and finally to enemies. The second approach involves cultivating equanimity by recognizing that all beings equally desire happiness and freedom from suffering. This leads to considering the advantages of selfless behavior and practicing the exchange of self-cherishing for cherishing others and wishing to protect them.

#### Emptiness^24^

This practice, considered a pure analytical meditation, focuses on the nature of phenomena, particularly the self, recognizing that the ordinary sense of self is based on a misapprehension. The practice begins by analyzing the way the sense of “me” appears to the mind. The core method is to logically question whether this “me” is identical with the body, the mind, the composite, or something separate from either. Emptiness is defined as the “unfindability”; of an object when logically searched for among its components. The goal of the logical pursuit is to arrive at this subtle appreciation of unfindability. This absence encountered upon searching is the phenomenon’s emptiness. The concept of emptiness is inextricably linked to the dependent nature of the object, which is called dependent origination. This understanding clarifies that the perception of a flower exists only in relation to the parts that make it up, but when searching for the flower among its parts, you will not find it. From this deep understanding of dependent origination ensues a rejection of any idea of intrinsic or inherent existence. The practice culminates by focusing the mind on this non-findability, thereby dissolving the instinctive clinging to a solid, inherent self. To deepen this realization of emptiness, the meditator must join the analysis with single-pointed concentration. This involves focusing the mind on the analysis by means of the previously acquired single-pointed concentration.

### Data Acquisition

#### Peripheral signals

##### Electrodermal activity – EDA

The EDA signal was acquired using an Empatica EmbracePlus health monitoring device. EmbracePlus is a wearable monitoring device that collects, processes, stores, and wirelessly transmit physiological parameters to a companion device (e.g., a smartphone) through a dedicated Care Lab App. The App collects data recorded by the EmbracePlus via Bluetooth® and securely uploads them to the Empatica Cloud using the smartphone’s internet connection. Raw data and digital biomarkers are subsequently accessed from the cloud via an AWS S3 Bucket. Among the several signals collected by the EmbracePlus (i.e., photoplethysmography, EDA, temperature, acceleration), we focused on the EDA as a correlate of sympathetic nervous system activity. The EDA signal was sampled at 4 Hz.

##### Electrocardiogram – ECG

The ECG signal was acquired with the peripheral physiological measurement module of a BE PLUS LTM from EBNeuro S.p.A., Firenze, Italy. The recording was performed at a sampling rate of 512 Hz, using two electrodes positioned below the left clavicle and above the right leg. The reference electrode corresponded to the electrical reference of the EEG cap.

##### Respiration – RESP

The RESP signal was acquired using the same BE PLUS LTM module as the ECG. The recording was performed at 512 Hz, using a piezoelectric belt recording thorax movement.

#### Electroencephalogram

The EEG signal was acquired using a BE PLUS LTM device from EBNeuro S.p.A, Firenze, Italy. We used a 19-channel EEG cap with electrodes positioned according to the international 10-20 standard. Electrodes impedances were kept below 20Kohm using conductive gel. Two additional electrodes were placed in the left and right mastoids for possible subsequent re-referencing. All signals were acquired at a sampling rate of 512 Hz.

### Physiological data analysis and feature extraction

#### Autonomic nervous system correlates

We extracted several features from the EDA, ECG and RESP signals to monitor subjects’ changes in ANS activity (Figure 1).

The EDA signal reflects changes in the skin conductance induced by eccrine sweat gland activity, which is under the only control of the sympathetic branch of the ANS. Accordingly, the analysis of the EDA is a reliable way to infer SNS dynamics^25^. The EDA signal is typically modelled as the sum of two components that carry non-overlapping information: a tonic component (or skin conductance level, SLC), a slow-varying component with a spectrum below 0.05 Hz, and a phasic component (or skin conductance response, SCR), which reflects short-term stimulus evoked responses^22^. In our context, SCRs might occur due to distractions or any internal emotional reactions during meditation.

Here, we used the cvxEDA model to extract tonic and phasic components of the EDA signal, as well as sudomotor nerve activity (SMNA) generating phasic responses^26^. Specifically, the signal was visually inspected for artefact removal, with portions of large artefacts being excluded from the analysis. Subsequently, the signal was normalized to have zero mean and unit variance before applying the cvxEDA model for numerical stability.

Then, starting from the estimated tonic, phasic and SMNA, several features related to SNS activity were derived, namely: the mean and standard deviation of tonic and phasic (i.e., **Mean Tonic, Std Tonic, Mean Phasic, Std Phasic**), the sum of peaks of the SMNA (i.e., **AmpSum**), and the power spectrum in the (0.045-0.25)Hz (i.e., **EDASYMP**^27^). More specifically, tonic-related features were estimated in consecutive windows of 60s, while phasic and SMNA features were estimated in consecutive windows of 10s.

The ECG signal was analyzed with NeuroKit2^28^ to extract the HRV. Among the several algorithms implemented in NeuroKit2, we used the Pan-Tompkins algorithm^29^ to extract the RR interval series (or NN series) from the ECG signal. Peak-detection artefacts were visually inspected and corrected using a cubic spline interpolation method. The obtained RR series were resampled to 4Hz to derive the HRV signal^21^. Subsequently, starting from the HRV signal we extracted several features in time, frequency and nonlinear domains, which are representative of ANS dynamics^30^. Unlike EDA, HRV is influenced by both SNS and PNS activity. Key time-domain indices included the mean and standard deviation of NN intervals (i.e., **MeanNN** and **SDNN**). We also computed the root-mean-square of successive differences (**RMSSD**) and the proportion of consecutive differences greater than 50 ms (**pNN50**). These indices quantify the amount of beat-to-beat variability, with RMSSD and pNN50 reflecting high-frequency variability related to parasympathetic modulation. To extract frequency-domain features, we estimated the power spectral density (PSD) of the HRV signal using Welch method. We then computed the power in the high-frequency (**HF**) band (0.15-0.4 Hz), reflecting parasympathetic activity, and low-frequency (**LF**) band (0.04-0.15 Hz), which reflects a mix of sympathetic and parasympathetic influences, and their ratio (**LF/HF**). Finally, we assessed the standard deviation of short-term interval variability (**SD1**) and long-term variability (**SD2**) via Poincaré plot. The **SD1/SD2** ratio was also calculated, as it can reflect the randomness/unpredictability of the heart rhythm and has been linked to autonomic balance. Features were estimated in 60s long windows.

The RESP signal was finally analyzed to extract the average breathing rate (**BR**) during each session.

#### Central nervous system correlates

The EEG signal was analyzed in terms of global power and connectivity metrics with EEGLAB^31^ and MATLAB Custom scripts.

First, we pre-processed EEG signals using a pipeline comprising

1. High-pass filtering (filter cutoff at 1Hz)
2. Notch filtering to remove line noise
3. Bad channel removal based on correlation criterion^32^
4. ICA decomposition^33^
5. Visual inspection for the identification and removal of artefactual components (e.g., eye, muscle). The EEG signal was then reconstructed using only components related to brain activity^34^.

Features were estimated using a sliding-window approach with 1-min-long non-overlapping windows. Within each of these windows, for each electrode, we estimated the power spectral density (PSD) using the Welch method with 5s-long consecutive windows, with 50% overlap, and a zero-padding equal to 2048 samples. A global measure of PSD was subsequently obtained by averaging the individual electrode PSDs across the entire scalp. Finally, the average PSD was then calculated for each session and expressed in decibels (dB).

In addition to spectral analysis, we assessed the dynamics of long-range neuronal organization through global functional connectivity metrics^35^. These metrics highlight the temporal synchronization between different cortical areas, providing complementary information to PSD measures. We computed two global connectivity metrics: (i) global magnitude of squared coherence (Coh)^35^ and (ii) global phase lag index (PLI)^36^. Global magnitude of squared coherence estimates the average linear consistency of phase differences between all pairs of electrodes. Global phase locking index, instead, measured the consistency of phase differences, being less sensitive to volume conduction issues compared to standard coherence.

Both power and connectivity metrics were calculated for the canonical EEG frequency bands: δ (1–4 Hz), θ (4–8 Hz), α (8–12 Hz), β (13–30 Hz), and γ (30–45 Hz).

### Statistical Analysis

To assess the effect of the experimental condition on each extracted feature, the Linear Mixed Model (LMM) of equation 1 was employed^37^.

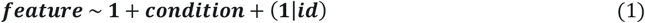

Particularly, we took advantage of including the subject as a random intercept, partitioning the total variance into between-subject variability and within-subject (residual) variability. This approach leads to more robust and precise estimates of the fixed effect (condition) compared to methods that treat subject-level variability as fixed. Furthermore, LMMs offer several key statistical advantages: they allow for the direct inclusion of all available data, effectively managing unbalanced or incomplete datasets (e.g., missing observations in specific sessions) without requiring imputation or subject exclusion.

The inclusion of the random intercept was justified using a Likelihood Ratio Test (LRT), comparing the full model against a fixed-effects-only model. Moreover, since we were fitting a model for each different feature, multiple hypothesis testing was controlled using the false-discovery-rate correction^38^. For those features that resulted in a significant model after FDR correction, post-hoc tests were performed using contrast-based t-tests on the fixed effects, followed by FDR correction.

### Data and Code Availability

Data and code leading to these results are available from the corresponding author upon reasonable request.

## Results

### Summary of findings

In Table 1 and 2, we report the summary results for the ANS and EEG features, respectively. All models were significant for the inclusion of the random intercept.

**Table 1.** Summary of statistics on ANS features.

|  | Feature | Model significance: p-value (FDR BKY) |
| --- | --- | --- |
| HRV | HF<sub>n</sub> | 0.360 (0) |
|  | LF/HF | 0.671 (0) |
|  | LF<sub>n</sub> | 0.393 (0) |
|  | MeanNN | **0.012 (1)** |
|  | RMSSD | **0.030 (1)** |
|  | SD1 | **0.030 (1)** |
|  | SD2 | **1.009E-06 (1)** |
|  | SDNN | **3.523E-05 (1)** |
|  | pNN50 | 0.726 (0) |
| EDA | Phasic Mean | **0.002 (1)** |
|  | Phasic Std | **0.002 (1)** |
|  | Ampsum | **0.002 (1)** |
|  | Tonic Mean | **4.166E-15 (1)** |
|  | Tonic Std | **0.001 (1)** |
|  | EDASYMP | **0.0301 (1)** |
| RESP | BR | **0.001 (1)** |

**Table 2.** Summary of statistics on EEG features.

|  | Feature | Model significance: p-value (FDR BKY) |
| --- | --- | --- |
| PSD | PSD<sub>δ</sub> | 0,140 (0) |
|  | PSD<sub>θ</sub> | **0,012 (1)** |
|  | PSD<sub>α</sub> | **1,770E-28 (1)** |
|  | PSD<sub>β</sub> | 0,185 (0) |
|  | PSD<sub>γ</sub> | **2,931E-13 (1)** |
| Coherence | Coh<sub>δ</sub> | 0,159 (0) |
|  | Coh<sub>θ</sub> | 0,326 (0) |
|  | Coh<sub>α</sub> | **6,227E-26 (1)** |
|  | Coh<sub>β</sub> | **0,0167 (1)** |
|  | Coh<sub>γ</sub> | 0,2948 (0) |
| PLI | PLI<sub>δ</sub> | **0,003 (1)** |
|  | PLI<sub>θ</sub> | 0,164 (0) |
|  | PLI<sub>α</sub> | **1,221E-20 (1)** |
|  | PLI<sub>β</sub> | **5,509E-08 (1)** |
|  | PLI<sub>γ</sub> | 0,848 (0) |

### ANS correlates

In Figure 3, we report the rainbow plots for the HRV features whose LMMs showed significant effects, along with the corresponding post hoc analyses. We observed a significant increase in **MeanNN** from the pre–EC rest to the post–EC2 condition. This increase in **MeanNN** indicates a slower heart rate, reflecting enhanced parasympathetic tone and a vagal response. This interpretation is further supported by the concurrent increases in **SDNN** and **SD2**, two indices of long-term vagal variability, consistent with elevated parasympathetic activity and greater overall autonomic flexibility. Moreover, **SDNN** and **SD2** displayed a more fine-grained temporal pattern, showing a gradual increase over the session and exhibiting distinct differences across the various meditation types.

**Figure 3.**
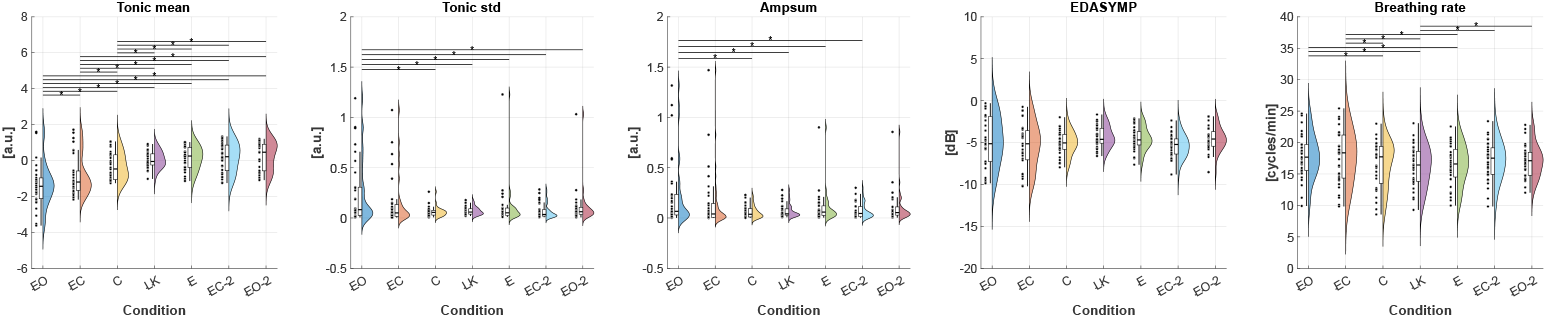
EDA and RESP changes. Post-hoc comparisons that were significant (p<0.05) are marked with an asterisk *.

In Figure 4, we report the rainbow plots for the EDA and RESP features whose LMMs showed significant effects, along with the corresponding post hoc analyses. We observed a gradual, fine-grained increase in the **Mean Tonic** component, indicative of heightened sympathetic arousal over the course of the session. This is further supported by the concurrent changes in **EDASYMP**, an index of sympathetic drive. However, post hoc analyses did not reveal significant pairwise differences for **EDASYMP**. Finally, we observed a significant decrease in **BR** during the meditation sessions compared to the EO and EC resting phases.

**Figure 4.**
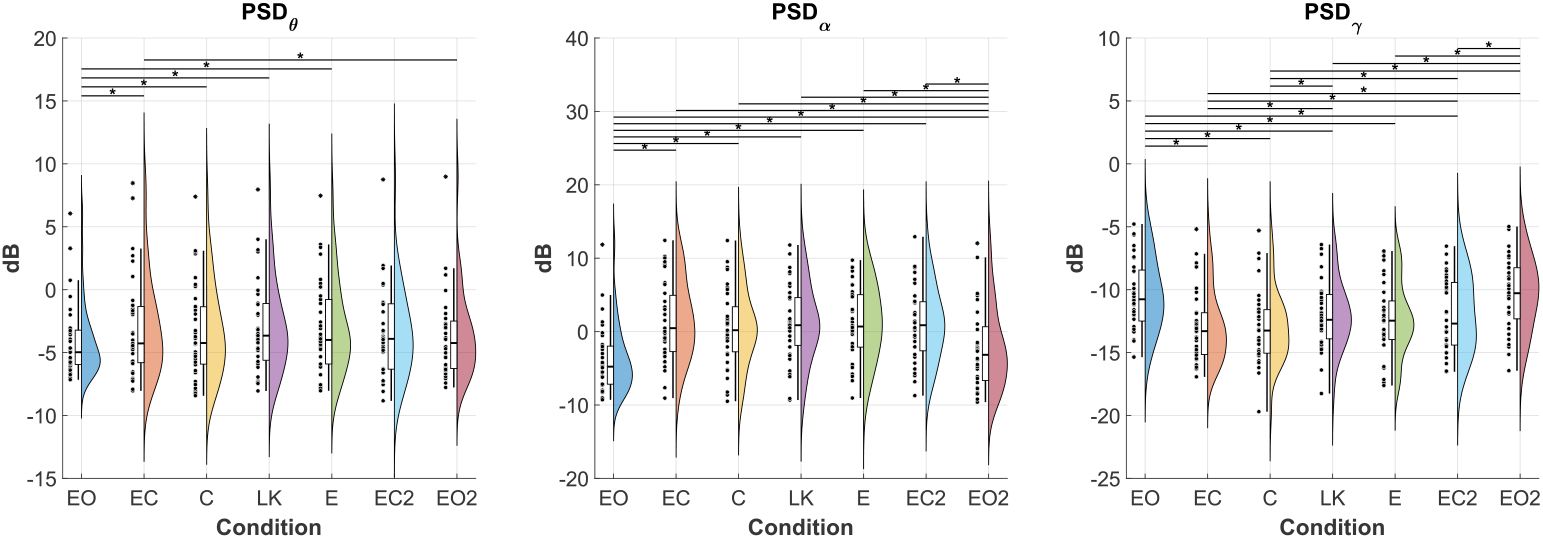
EEG PSD changes. Post-hoc comparisons that were significant (p<0.05) are marked with an asterisk *.

### EEG correlates

In Figure 5, we report the rainbow plots for the EEG power spectral density (PSD) features whose LMMs showed significant effects, along with the corresponding post hoc analyses. The main differences were observed in the gamma band, where a gradual increase in PSD was evident over the course of the experiment. This increase was also significant between the concentrative session (C) and the two analytical sessions (LK and E). Additionally, we observed an increase in both theta and alpha activity during both the eyes-closed (EC and EC2) and meditation sessions (C, LK, E) compared to the eyes-open conditions (EO and EO2).

**Figure 5.**
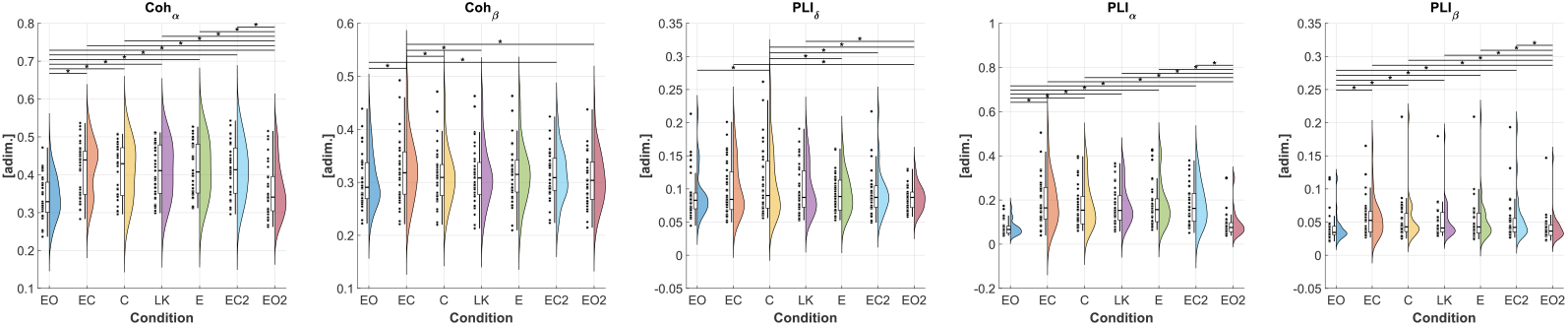
EEG functional connectivity changes. Post-hoc comparisons that were significant (p<0.05) are marked with an asterisk *.

In Figure 6, we report the rainbow plots for the EEG functional connectivity features whose LMMs showed significant effects, along with the corresponding post hoc analyses. The main changes were driven by differences between the eyes-open conditions (EO and EO2) and both the eyes-closed (EC and EC2) and meditation sessions (C, LK, E) in the alpha band for both Coherence and PLI, and in the beta band for PLI. Interestingly, we observed Specifically, we observed a significant difference in PLI in in delta band between C and E.

## Discussion

This study analyzed the central and peripheral neural correlates of concentrative and analytical forms of meditation in a group of experienced Tibetan monks meditators. Thirty-five experienced Tibetan monks from the Sera Jey Monastic Institute, all with over 10 years of formal monastic education in the same cultural and social context, participated in a within-subjects experimental protocol comprising Concentrative, Loving-Kindness, and Emptiness meditation, the latter two representing two distinct forms of analytical practice. We showed that the three meditation types resulted in differential changes to both autonomic correlates (EDA, HRV, and respiration), and central neural correlates (EEG), and that these changes, altogether, converge upon the delineation of a relaxed-alertness state, which is a primary outcome of meditation and strongly associated with enhanced well-being^39^.

### Gamma power increase: augmented cognitive processing during analytical meditation

The results showed higher levels of gamma-band power in the two EO conditions than in all the other experimental conditions (EC, C, LK and EC-2), which were all conducted with the eyes closed. This finding is consistent with previous observations indicating that eye closure causes focal decreases in gamma power (30–100 Hz) over the occipital cortex and supports the hypothesis that higher gamma power reflects the active processing of visual information^40^.

No gamma power differences were found between EC rest and Concentrative meditation (C), which is suggestive of the lack of a state effect. This finding contrasts with previous evidence, which showed gamma-band changes during Concentrative meditation, typically emerging after 20–30 minutes of sustained practice^11^. The shorter duration employed in our protocol (15 minutes) may therefore have limited the likelihood of detecting such effects.

Gamma power in LK meditation was higher than in EC rest. In this light, increased gamma activity has been previously observed in meditation practitioners during open monitoring (OM) Vipassana meditation with respect to mind-wandering ^41^, and during OM mindfulness meditation compared to resting state^42^. However, the research conducted by Braboszcz and colleagues^43^, who examined the potential increases in parieto-occipital and frontal gamma-band power in three distinct meditation traditions (Vipassana, Himalayan Yoga and Isha Shoonya) compared to a control group - during both meditative state and instructed mind-wandering – reported no substantial effects.

It has been proposed that EEG gamma-band activity is the neural correlate of various cognitive functions. These include the ongoing stream and contents of consciousness^44^, long-range neuronal communication^45,46^ and attention^47,48^. This suggests that an increase in EEG gamma-band activity may provide a background for periods of greater cognitive engagement, such as those experienced during some meditative practices in which this engagement is present, particularly analytical ones^11,24,49^.

In fact, the three meditations carried out during the experimental session belonged to three different families^50^, each involving specific mental processes. Concentrative meditation falls within the Focused Attention (FA) family; Loving Kindness meditation to the constructive family which also includes compassion meditation; Emptiness meditation to the deconstructive family including different sorts of insight meditation. Both LK and E are classified as analytical meditations, as they rely on reflective and conceptual processes to transform perception and understanding.

Considering the profound cognitive engagement common to all three forms of meditation, we can hypothesize that the greater gamma power during LK meditation may represent a greater emotional involvement compared to C and E meditation. This difference is related to the specific demands of each practice. *Concentrative* meditation involves maintaining attention on a single object, such as the breath, while monitoring and stabilizing awareness^51^. *Loving-kindness* meditation, although aimed at cultivating a mental state that is emotionally characterized by unconditional compassionate love, achieves this through substantial cognitive engagement. This reflective component introduces an analytical element absent in purely concentrative practices. *Emptiness* Meditation, by contrast, seeks to deconstruct the illusion of how things superficially appear to exist and entailing the inquiry into the non-existence of an inherent, permanent self. It necessitates a profound, step-by-step examination of objects and concepts to comprehend their lack of inherent existence, therefore it can be considered a pure analytical meditation as opposed to a straightforward cessation of thought typical of concentrative meditation.

### Increased vagal tone and HRV changes

Analysis of meanNN and HRV features indicated a generalized slowdown in heart rate across the experimental sessions, indicative of enhanced parasympathetic vagal modulation. Concurrently, increased overall (both parasympathetic and sympathetic) and long-term HRV was observed, as evidenced by alterations in SDNN and SD2 over the session, which suggests an increase in overall autonomic flexibility. The gradual increase of these features, with distinct differences observed across the various meditation types, indicates the possible fine modulation of autonomic control to allow better adaptation of heart activity to the different meditation types.

It is important to note that, irrespective of the phase of the experimental session (be it rest or meditation), the meditators demonstrate remarkably elevated levels of vagal tone. This is substantiated by the augmented levels of RMSSD, an HRV feature in the time domain that is widely regarded as a reliable indicator of vagal tone^52^. A body of experimental evidence has demonstrated that elevated vagal tone is associated with enhanced emotional regulation and health, as well as better executive functions, including mental flexibility, working memory, attention regulation and mental imagery^53–56^. Therefore, it may be hypothesized that elevated vagal tone is a trait of experienced meditators that has been strengthened through protracted meditation practice, which is essential for optimal meditation but concomitantly results in enhanced cognitive abilities and well-being.

### Breathing frequency regulation

The study found a reduced breathing rate during all three meditation types compared to both EO and EC rest, with no significant differences among the meditation conditions. A decrease in respiration rate during meditation relative to the initial baseline is expected. Many meditation techniques, in fact, involve focusing on the breath and deliberately slowing it down. This often naturally results in a slower, more regular breathing rhythm, which promotes relaxation and a sense of calm. A particularly pronounced deceleration in the rate of respiration has been previously documented in long-term meditators during periods of meditation, with a more marked slowdown observed during more intensive practices, such as retreats. A slower respiration rate has also been found in experienced meditators during rest periods (see^57^ for a review). Breathing can significantly affect heart rate variability (HRV), and controlled breathing at slower rates effectively promotes autonomic function and optimizes HRV^58^.

### Arousal and relaxed vigilance

Analysis of electrodermal activity (EDA) revealed a progressive increase in mean tonic EDA, i.e. the slow, background level of skin conductance-over the initial stage of the experimental session (from EO to LK). The maximum mean tonic value achieved during LK meditation was, then, sustained throughout E meditation and the subsequent rest periods (EC-2 and EO-2). Tonic EDA is widely regarded as an indicator of tonic sympathetic activity, reflecting sustained autonomic arousal and vigilance over time. Changes in tonic EDA levels can serve as biomarkers to assess an individual’s overall physiological and psychological arousal state, which in turn can influence cognitive functions such as attention, memory, and executive control.

In conventional Buddhist texts, meditation is a condition of “relaxed alertness”, i.e. a mental state that is characterized by maintaining balance between hypoarousal (excessive relaxation and drowsiness) and hyperarousal (agitation, restlessness)^59,60^. This state is foundational to proceed with specific meditation practices aimed at reaching different targets. Nonetheless, most research conducted on meditation has only emphasized its calming effects and its capacity to reduce arousal, while paying less attention to its potential to enhance alertness.

The current results, which indicate an increase in tonic sympathetic activity along with a slight increase in vagal modulation, highlight the ability of experienced meditation practitioners to achieve, through the meditation sessions, a state of “awareness” which, in meditation, refers precisely to the attainment of a state of alert relaxation. This is supported by the observation that the highest mean tonic EDA value is reached during LK meditation and maintained for the rest of the session.

The increased tonic EDA during LK and E meditations, which is associated with higher gamma-band activity, supports the hypothesis that meditation practices requiring heightened cognitive engagement necessitate an increased level of overall physiological and psychological arousal.

The achievement of alert relaxation through meditation is supported by a number of neuroimaging studies indicating that meditation-related cerebral activation corresponds to tonic alertness–related brain areas (lateral PFC, inferior parietal, and anterior insula)^51,61,62^. In addition, an increased level of connectivity was identified between the anterior and posterior nodes of the Default Mode Network (DMN) in experienced meditators during meditation. This connectivity is known to decrease during slow-wave sleep and vegetative states, and an increase in this connectivity is likely to be indicative of increased wakefulness^63,64^. The finding of increased intra-DMN connectivity in expert meditators, even during rest periods, may be indicative of neuroplastic changes that underpin meditative skills^65^.

The present body of evidence supporting the ability to achieve a state of alert relaxation through meditation practice also supports the broader concept of autonomic control in terms of autonomic flexibility as opposed to autonomic balance. Indeed, it is evident that the two branches of the autonomic nervous system can collaborate adaptively to optimize the body’s psychophysical functioning, rather than necessarily working in antagonism. Furthermore, it is worth noting that this autonomic flexibility can be developed through mental training.

### Other observed changes: Theta and alpha power and global connectivity metrics

Lower alpha and theta activity were only observed during the eyes-open conditions (EO and EO2) compared to the eyes-closed conditions (EC and EC2) and meditation sessions (C, LK and E). However no differences were observed between different types of mediation. Although we previously noted large differences in theta and alpha power in an earlier study in different meditation typologies ^11^, significant effects of this kind typically emerge after the first 20 minutes of concentrative meditation. This may therefore explain the discrepancy with the present results, which are limited to 15 minutes of concentrative practice. In this light, the short duration of the concentrative session may have contributed to findings consistent with previous reports suggesting that increases in alpha power are not generally considered a reliable indicator of meditative states^41^. In this context the duration of the meditation session may play a key role. Moreover, an investigation into a group of Vipasana meditation practitioners revealed that there were no significant differences between the 7-11 Hz alpha band power during meditation and during a mind-wandering control task^43^. In this light, we have previously observed that higher alpha amplitude is more indicative of expert meditators engaged in long-term sessions of concentrative type^11^. Increases in alpha power have been identified in tasks requiring the redirection of attention towards an internal object^66^. Furthermore, modulation of the alpha rhythm has been hypothesized to play a role in selective top-down attention processes by regulating thalamo-cortical sensory transmission and thereby contributing to functional inhibition^67,68^. Because of these premises, it can be argued that changes in trait alpha amplitude in expert meditators may be attributable to their extensive meditation training, which may have enhanced their capacity to internalize focus and to increase gating of distracting stimuli.

Earlier EEG studies have reported meditation-related increases in the theta band^11,69^. Nevertheless, subsequent research findings have highlighted that increases in theta activity are not widespread over the whole scalp but localized to the frontal midline regions of meditators^70–72^. This phenomenon is manifested predominantly during concentrative meditation and is attributable to medial prefrontal cortex activity^73^. As with the alpha power, it is conceivable that the increased theta power observed during meditation and eyes-closed conditions could be interpreted as an indication of an experienced meditator’s characteristic trait.

Global Connectivity features did not show specific patterns across conditions, suggesting that more fine-grained analysis focusing on local features may be a more practicable avenue to identify connectivity changes related to different meditative states. However, pursuing this path presents a significant methodological challenge: including numerous local connectivity metrics would substantially increase the number of features tested, thereby lowering the statistical power and inflating the risk of Type I error (false positives) due to multiple comparisons.

To mitigate this issue, confirmatory modeling approaches (e.g., Dynamic Causal Models^74^) would offer a more robust solution. Here, since our study was primarily exploratory in nature, we chose to focus on global metrics. Among the global findings, the most relevant change is the increase in global PLI in the delta band during *Emptiness* compared to *Concentrative*. This finding could be interpreted alongside the previously noted increase in global PSD in the gamma band, suggesting a potential link to increased working memory and enhanced internally focused processing during the Emptiness state^51^.

### Limitations and future development

The fixed order of experimental conditions (C→LK→E) represents a potential confounding factor in our analysis. This design choice prevents us from fully disentangling the effect of each specific session from potential order effects, such as fatigue or cumulative carry-over effects. However, this fixed sequence was structurally required by the established methodology of these specific contemplative practices inside the Tibetan tradition. Concentrative meditation is typically used to induce a foundational state of calm and mental stabilization, which is considered mandatory for effectively engaging in the successive, more advanced stages of meditation. Furthermore, the placement of the *Loving-Kindness* session before the *Emptiness* session is also grounded in practice: the emotional and perceptual nature of *Loving-Kindness* is commonly sequenced prior to the highly conceptual and abstract nature of the *Emptiness* practice. Therefore, while we acknowledge the methodological limitation of not employing a counterbalanced design, the chosen fixed order reflects the ecological validity and structural integrity required for the authentic progression through these meditative states.

A possible workaround for this limitation would be to acquire an equally long resting-state session for each participant, at the beginning or end of the overall experiment. While each participant completed three fixed-order meditation sessions (C, LK, E), this additional RS session would provide a crucial control for possible time-on-task or non-specific time effects that might otherwise be confounded with the order effects inherent to the meditative practice itself. Future work could consider incorporating this control practice.

### Future development

The multivariate acquisition of concomitant peripheral physiological signals (representative of autonomic nervous system dynamics) alongside central brain activity and connectivity measures estimated from EEG, provided a rich overview of the mind-body changes induced by meditation. In this light, we suggest that multivariate approaches are preferable, when possible, to better characterize and interpret the effects of different mental states on bodily changes, thereby promoting structured methodologies for the study of non-ordinary states of consciousness^75^.

Future studies may benefit from considering additional complementary features to further characterize the associated bodily changes. Furthermore, the use of confirmatory modeling techniques could be explicitly tested to delineate the relationships and dependencies between central and peripheral measures, moving toward a more comprehensive analysis of the whole-system dynamics of mind-body interactions.

### Declaration of generative AI and AI-assisted technologies in the writing process

During the preparation of this work the authors used ChatGPT 5.1 and GEMINI Flash 2.5 in order to improve the readability and language of the manuscript. After using these tools/services, the authors reviewed and edited the content as needed and take full responsibility for the content of the published article.

## Acknowledgments

The research leading to these results received partial funding from the Italian Ministry of Education and Research (MIUR) in the framework of the ForeLab Project (Departments of Excellence). This research is partly funded by the European Union – Next Generation EU, in the context of The National Recovery and Resilience Plan, Investment 1.5 Ecosystems of Innovation, Project Tuscany Health Ecosystem (THE), Spoke 3 “Advanced technologies, methods, materials and health analytics” CUP: I 3C22000780001.

## Author contributions

All authors designed the research. A. L. C., J. TH., J. S., J. TS., and B. N., collected the data. A. L. C., M. S. and A. G., analyzed the data. A. L. C., M. A., L. S., and B. N interpreted the results and drafted the article. All authors wrote the article.

### Declaration of interest

Authors declare no competing interests.

## Notes

### Competing Interest Statement

The authors have declared no competing interest.

